# Pipette, mix, repeat: A reduced-order fluid model and protocol survey to evaluate 96-well mixing protocols

**DOI:** 10.64898/2025.12.27.696695

**Authors:** Michael R. King

## Abstract

Mixing in 96-well plates is a fundamental operation in biology and bioengineering, yet the most common strategy, pipetting up and down to “mix well”, is rarely quantified or justified. Rules of thumb such as “three mixing cycles” coexist with protocols that recommend 5, 10, or 20 pipette strokes per step, and there is limited fluid-mechanical guidance on when these choices are adequate. Here, a reduced-order compartment model of advection-diffusion in a single 96-well plate well is developed, in which the fluid is discretized into hundreds of macrocells with finite-volume-style neighbor exchange. A pipette jet is represented as a localized enhancement of exchange along a pipette-aligned “jet footprint,” parameterized by an effective jet strength, effective jet footprint size as a proxy for cycled volume fraction, pipette angle, and centered versus off-center tip placement. Repeated aspirate-dispense cycles are simulated and mixing quantified by the decay of normalized concentration variance. The reduced model qualitatively reproduces the mixing hierarchy reported in high-fidelity COMSOL simulations of repetitive pipetting, with higher jet strength and jet footprint substantially accelerating homogenization, while gentle, small-footprint jets often require more than three cycles to reach near-uniform thresholds. Systematic variation of pipette angle and tip position shows that moderately off-center, moderately tilted pipetting modestly improves mixing relative to strictly vertical, centered configurations, with angle and tip position effects remaining small compared to jet strength and footprint size. In parallel, a survey of 96-well pipette mixing instructions in online research forums, commercial assay protocols, and published papers was carried out, extracting explicit “pipetted up and down N times” stroke counts. The resulting distribution varies from 2 to 20 strokes per mixing step, with no clear consensus. This combined modeling and survey analysis indicates that three cycles are not sufficient for many instances of 96-well pipette mixing, and provides a fast, interpretable tool for rationalizing and redesigning pipette-mixing protocols in high-throughput experiments.

## Introduction

Microtiter plates are a workhorse of modern biology, biotechnology, and drug discovery, supporting high-throughput screening, enzyme kinetics, cell culture, and diagnostic assays in standardized 96-, 384-, and 1536-well formats [1,2]. The 96-well plate represents a key compromise between throughput, volume, and compatibility with manual or semi-automated workflows, and has therefore become deeply embedded in both academic and industrial practice. As methods have miniaturized, attention has turned to the fluid dynamics of microplate wells, including gas–liquid mass transfer, shear stress, and mixing times, because these physical processes can strongly affect cell physiology, reaction kinetics, and assay readouts [1–4].

A rich literature has characterized mixing and hydrodynamics in shaken microtiter plates and related small-scale vessels [5]. Studies have quantified the dependence of mixing time and power input on shaking diameter, frequency, and filling volume [3,4,6], analyzed flow patterns and shear stress distributions in culture well plates [7], and developed CFD-based or empirical models to support scale-up and clone selection [4,8]. Engineering characterization of individual wells from 24- and 96-well plates has highlighted the sensitivity of flow regimes to geometry and operating conditions [9]. These works collectively provide a detailed picture of mixing under orbital shaking and related agitation strategies, and they have informed guidelines for using plates as “mini-bioreactors” [1,2,4,10,11].

By contrast, pipette-driven mixing in 96-well plates—aspirating and dispensing liquid with a pipette tip directly in each well—has received relatively little formal modeling attention. Weiss and co-workers pioneered a compartment model of mixing in 96-well plates, combining experimental measurements with soluble pH indicators and immobilized fluorescence sensors [8]. Their model used a limited number of mixed elements to describe how small additions of alkali dispersed under various addition methods (multichannel pipette, piston pump, piezoelectric actuator). Subsequent studies have advanced CFD simulations and hydrodynamic models of microtiter wells [4,7–9,12,13], but these generally focus on shaken or stirred configurations rather than repetitive aspiration–dispense cycles with a pipette tip. In parallel, pharmacokinetic and permeability studies have demonstrated that inadequate mixing in 96-well sandwich plates can bias measured permeabilities, and that agitation method (orbital shaking vs magnetic stirring) has an effect [14].

Despite this growing theoretical and experimental literature, day-to-day microplate practice still relies heavily on simple verbal instructions such as “mix well” or “pipette up and down to mix.” Informal rules of thumb—for example, that “three mixing cycles” are sufficient—are widely repeated but rarely justified with fluid-mechanical reasoning. Vendor application notes and kit instructions sometimes specify stroke counts (e.g., “mix 10 times”), but there is no consensus and little connection to underlying fluid physics [12,15]. In addition, many protocols omit stroke counts entirely, leaving users to decide how much mixing effort to expend, and making it difficult to assess when under- or over-mixing might compromise data quality.

The present work addresses this gap from two complementary angles. First, a reduced-order fluid model for pipette mixing in a single well of a 96-well plate is developed. Inspired by finite-volume advection–diffusion formulations and earlier compartment models [8,9], the well volume is discretized into a finite grid of macrocells, with neighbor-to-neighbor exchange representing combined advection and diffusion. A pipette jet was represented as a localized enhancement of exchange along a “jet footprint” oriented with the pipette axis, with parameters for jet strength, effective jet footprint size as a proxy for cycled volume fraction, pipette angle, and centered versus off-center tip placement. This yielded a computationally inexpensive model that preserves mass conservation and allows rapid parameter sweeps over stroke count, geometry, and mixing aggressiveness. The model was qualitatively validated by comparing predicted mixing trends to representative CFD and experimental studies of mixing in microtiter plates [3,4,6,9,12,13].

Second, a survey of 96-well pipette-mixing instructions was carried out, spanning the r/labrats subreddit of Reddit.com, commercial kit manuals that explicitly specify the number of “pipetted up and down *N* times” strokes, and a small number of peer-reviewed articles. This allows one to construct an empirical distribution of the number of pipette strokes used in contemporary practice and to compare it directly to the mixing behavior predicted by the reduced-order model.

The present goals were (i) to test, in a controlled modeling framework, whether there is a clear number of mixing cycles that emerges as sufficient under realistic parameter choices; (ii) to explore how pipette angle and tip placement modulate mixing efficiency; and (iii) to place these predictions in the context of actual laboratory practice documented in online forums and documents. In doing so, a GPT-native workflow in computational biology is illustrated: a large language model was used to help scaffold model code, prototype figures, and draft text, while all equations, simulations, and interpretations were curated and owned by the human author, in line with emerging guidance on responsible use of generative AI in computational biology [16–21].

## Methods

### Overview

A reduced-order model of solute mixing in a single well of a 96-well plate during repeated pipette aspiration–dispense cycles was constructed. The well volume was discretized into a finite grid of “macrocells” with assigned volumes and neighbor connections. Concentration transport is represented as a linear update rule describing exchange of solute between neighboring macrocells, with coefficients representing combined advection and diffusion. A pipette jet was modeled as a time-dependent enhancement of exchange within a subset of macrocells defining a “jet footprint,” oriented along the pipette axis. The model was implemented in Python (**Appendix**) and iterated numerically to simulate sequences of mixing cycles under varied geometric and protocol parameters.

In parallel, a survey of online protocol descriptions was carried out to extract the number of pipette strokes used for mixing in 96-well plates and these were compiled into a dataset for comparison with model predictions.

### Well geometry and macrocell discretization

A single round well was considered, representative of a standard 96-well plate. As a starting point, typical dimensions were adopted from plate specifications and prior engineering characterizations: well diameter ≈6.4 mm, and working fluid height ≈8–10 mm for volumes of 200–250 µL [2,9]. The fluid volume was approximated as a right circular cylinder and discretized into a regular Cartesian grid of macrocells representing the horizontal cross-section. In most simulations, a 16 × 16 grid (256 macrocells) was used, balancing spatial resolution against computational cost, with vertical recirculation represented implicitly by a global mixing term.

Each macrocell 𝑖 has volume 𝑉*_i_* and concentration 𝑐*_i_*(𝑡). The fluid was assumed incompressible and density-matched across macrocells. Neighboring macrocells (sharing a face) exchange solute according to effective exchange coefficients 𝑘*_ij_* (s^−1^) that represent the net effect of local advection and diffusion across the interface. In the absence of pipette action, the coefficients are isotropic and uniform:

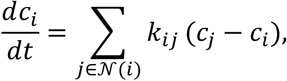

where *N*(*i*) is the set of neighboring macrocells sharing a face with macrocell 𝑖, and 𝑘*_ij_* = *k*_0_ for all neighboring pairs. This structure guarantees conservation of total solute mass and tends toward homogenization. In addition to nearest-neighbor exchange, a weak global recirculation term was included to relax each macrocell slightly toward the global mean concentration per cycle, with its strength increasing with jet strength and footprint size, to mimic bulk three-dimensional recirculating flows.

This formulation is conceptually similar to earlier compartment models of mixing in microtiter plates, which used a limited number of mixed zones to match experimental dye dispersion profiles [8,9], and to finite-volume approximations of advection–diffusion equations underlying CFD simulations of microtiter hydrodynamics [4,7,12,13].

### Representation of the pipette jet

To represent the pipette jet, a subset of macrocells was defined as belonging to the jet footprint—namely, those intersecting the notional jet core emanating from the pipette tip (**Figure 1**). The jet footprint was parameterized by:

- <u>Pipette angle</u> 𝜃 (0° = vertical, 30–60° = moderate tilt);
- <u>Tip position</u> in the horizontal plane (centered vs off-center toward the wall);
- <u>Footprint radius</u> 𝑟*_f_*, determining how many macrocells around the jet core are affected.

**Figure 1.**
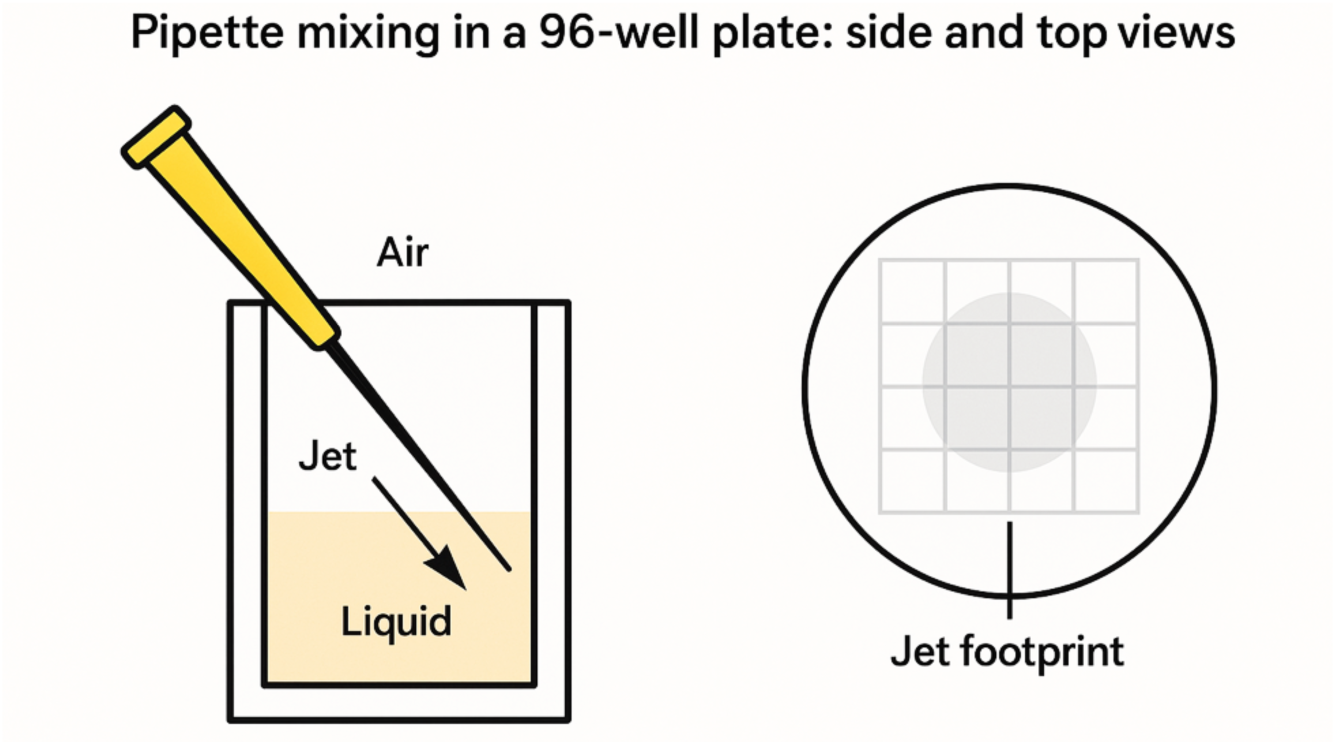
Schematic of the 96-well geometry, macrocell discretization, and jet footprint. Conceptual illustration of the reduced-order model. The well volume is discretized into a grid of macrocells (shown as a regular lattice), each exchanging mass with its four face neighbors. The pipette tip is positioned above the well at a specified angle and lateral offset, defining a pipette axis that penetrates the fluid. Macrocells lying within a specified distance of this axis and depth interval form the “jet footprint” (shaded region), in which neighbor exchange rates are enhanced during the dispense phase of each stroke. This construction approximates the advective mixing induced by the pipette jet while preserving a simple, finite-volume–style structure.

Within the footprint, the exchange coefficients 𝑘*_ij_* were increased during the dispense phase of each stroke to mimic the enhanced mixing driven by the jet. One may write

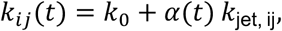

where 𝑘_jet, ij_ > 0 for edges within the footprint and 0 elsewhere, and 𝛼(𝑡) is a square-wave function that is 1 during dispense phases and 0 during aspiration and rest. The effective jet strength is controlled by a scalar parameter 𝐾_jet_, such that 𝑘_jet, ij_ scales proportionally to 𝐾_jet_. In practice, “gentle” jets (𝐾_jet_ comparable to 𝑘_0_) and “aggressive” jets (𝐾_jet_ ≫ 𝑘_0_) were explored.

A pipette stroke was defined conceptually by the cycled volume fraction ϕ, the fraction of total well volume aspirated and dispensed per stroke. Although the model does not explicitly remove and re-inject fluid parcels or track ϕ directly, the enhanced exchange within the footprint, combined with the global coupling of macrocells, approximates the net effect of repeatedly drawing fluid into the tip and ejecting it back along a jet aligned with the pipette axis. Jet footprint shapes and relative strengths were chosen to qualitatively reflect flow structures seen in CFD and experimental visualizations of jets and mixing in microwell systems [4,8,9,12,13].

### Dimensionless formulation and mixing metric

The time was nondimensionalized by a characteristic mixing time 1/𝑘_0_ and concentrations by the initial concentration difference between “labelled” and “unlabelled” macrocells. The ODE system can be written compactly as

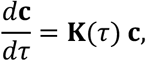

where 𝜏 = 𝑘_0_𝑡 is dimensionless time and 𝐊(𝜏) is the time-dependent rate matrix whose off-diagonal entries are 𝑘*_ij_* /𝑘_0_ and whose diagonal entries enforce conservation.

To quantify mixing, the normalized concentration variance was used:

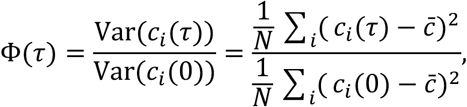

where 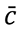 is the volume-weighted mean concentration and 𝑁 is the number of macrocells. A perfectly mixed state corresponds to Φ → 0. “Operationally mixed” thresholds were defined such as Φ = 0.05 or Φ = 0.01, inspired by tolerances used in mixing time studies [3,4,7,12].

### Simulation protocol and numerical integration

Simulations were implemented in Python using NumPy. Each aspirate–dispense cycle was represented as a single application of the linear mixing operator, which aggregates the effects of aspiration and dispense within that cycle. Because the exchange process is linear in concentrations, the absolute value of Δ𝜏_cycle_ can be interpreted as scaling the effective rate of mixing per stroke; relative changes in cycle count and jet parameters were explored rather than attempting to calibrate against an absolute physical time.

Four protocol scenarios were defined:

1. <u>Gentle pipetting, small footprint</u>: low 𝐾_jet_, small footprint radius, moderate 𝜙.
2. <u>Gentle pipetting, large footprint</u>: low 𝐾_jet_, larger footprint covering more of the well.
3. Aggressive pipetting, small footprint: high 𝐾_jet_, small footprint.
4. Aggressive pipetting, large footprint: high 𝐾_jet_, large footprint.

For each scenario, 1–12 cycles were simulated and Φ recorded after each full cycle. Parameter sweeps over pipette angle (0–60°), tip position (centered vs off-center), and footprint radius were also performed, and the resulting stroke counts required to reach Φ≤0.05 summarized.

To test sensitivity to grid resolution, selected simulations were repeated with coarser and finer macrocell grids (e.g., 8×8 and 24×24), confirming that qualitative trends in Φ vs cycle number and the relative ordering of scenarios were robust to grid refinement.

### Qualitative validation against existing microtiter mixing studies

Although this reduced-order model does not reproduce detailed velocity fields, it is closely related to finite-volume discretizations of advection–diffusion and to earlier compartment models used to fit intensity fields in 96-well plates [8,9]. Qualitative consistency with published mixing and hydrodynamics studies was therefore sought:

- Weiss et al. demonstrated that a multi-element mixing model can capture dispersion of small reagent additions in 96-well plates, and that mixing times depend strongly on addition method and geometry [8].
- Li et al. and Montes-Serrano et al. showed that mixing times and volumetric power input in shaken microwell systems decrease with increased agitation and that geometry strongly modulates mixing behavior [3,4].
- CFD-based 𝑘*_L_*a models in microtiter plates similarly emphasize the link between local flow features, energy dissipation, and global mass transfer [8,12,13].
- Eriksen et al. demonstrated that insufficient mixing in 96-well sandwich permeability assays leads to systematically biased permeabilities, and that agitation method has an effect [14].

To align the present model with these observations, parameter ranges were chosen such that:

- “Aggressive” jet scenarios achieved rapid drops in Φ over a few cycles, analogous to high-power mixing regimes.
- “Gentle” scenarios mixed more slowly and sometimes failed to reach Φ ≤ 0.05 even after many cycles, consistent with low-power or poorly configured agitation.
- Larger jet footprints were generally more efficient than smaller ones at fixed jet strength, reflecting the benefit of engaging more of the well volume per stroke.

A one-to-one calibration between Φ(cycle) and experimental mixing times was not attempted, but these qualitative correspondences used to avoid clearly unphysical parameter choices.

### Survey of pipette mixing stroke counts

To characterize current practice in 96-well pipette mixing, a search of recent protocols was conducted. The following sources were focused on:

- <u>Research forum posts</u> in the Reddit.com subreddit r/LabRats, and Researchgate.net.
- <u>Commercial protocols and application notes</u> from vendors of 96-well-based kits and mixers that specified stroke counts.
- <u>Protocol-centric journals</u>, including JoVE and Scientific Reports, where detailed, stepwise methods are routinely described;

Searches were performed using combinations of terms such as “96-well, “plate,” “mixing,” and “pipette” in full-text searches. From the resulting hits, protocols were manually extracted that contained explicit stroke counts for pipette mixing in 96-well wells (e.g., “pipette up and down 10 times”).

Records were included if they:

1. Used a 96-well plate as the primary vessel, and discussed pipetting as the mixing approach;
2. Provided an explicit number of strokes or cycles (integer counts such as 3, 5, 10, or 20), or in one case, “couple of times” was interpreted as 2.

These data were compiled into a table. In total, the curated dataset comprises 42 protocol instances, combining research forum posts, vendor documents, and peer-reviewed articles, each contributing at least one explicit pipette-mixing stroke count in a 96-well format. A histogram was plotted of the number of strokes per mixing step across all extracted protocols. Because the sample size, while reasonable, is not exhaustive and may be biased toward protocols that choose to mention explicit stroke counts, this analysis is descriptive and intended to provide context for the modeling results rather than a definitive census.

### Use of large language models

A large language model (ChatGPT, OpenAI; GPT-5.1 Thinking) was used as an assistive tool to prototype Python code for the reduced-order mixing model, suggest parameterizations, and draft preliminary text. All code, equations, and manuscript content were reviewed, edited, and validated by the author, who takes full responsibility for the final work.

## Results and Discussion

### Reduced-order model reproduces expected mixing hierarchies

**Figure 2** shows the normalized concentration variance Φ versus the number of pipette cycles for the four protocol scenarios. As expected, aggressive jets with large footprints (scenario 4) produce rapid mixing: Φ falls below 0.05 within roughly 3–4 cycles and below 0.01 within about 6–8 cycles. Aggressive jets with small footprints (scenario 3) mix somewhat more slowly, typically requiring a few additional cycles to achieve the same thresholds.

**Figure 2.**
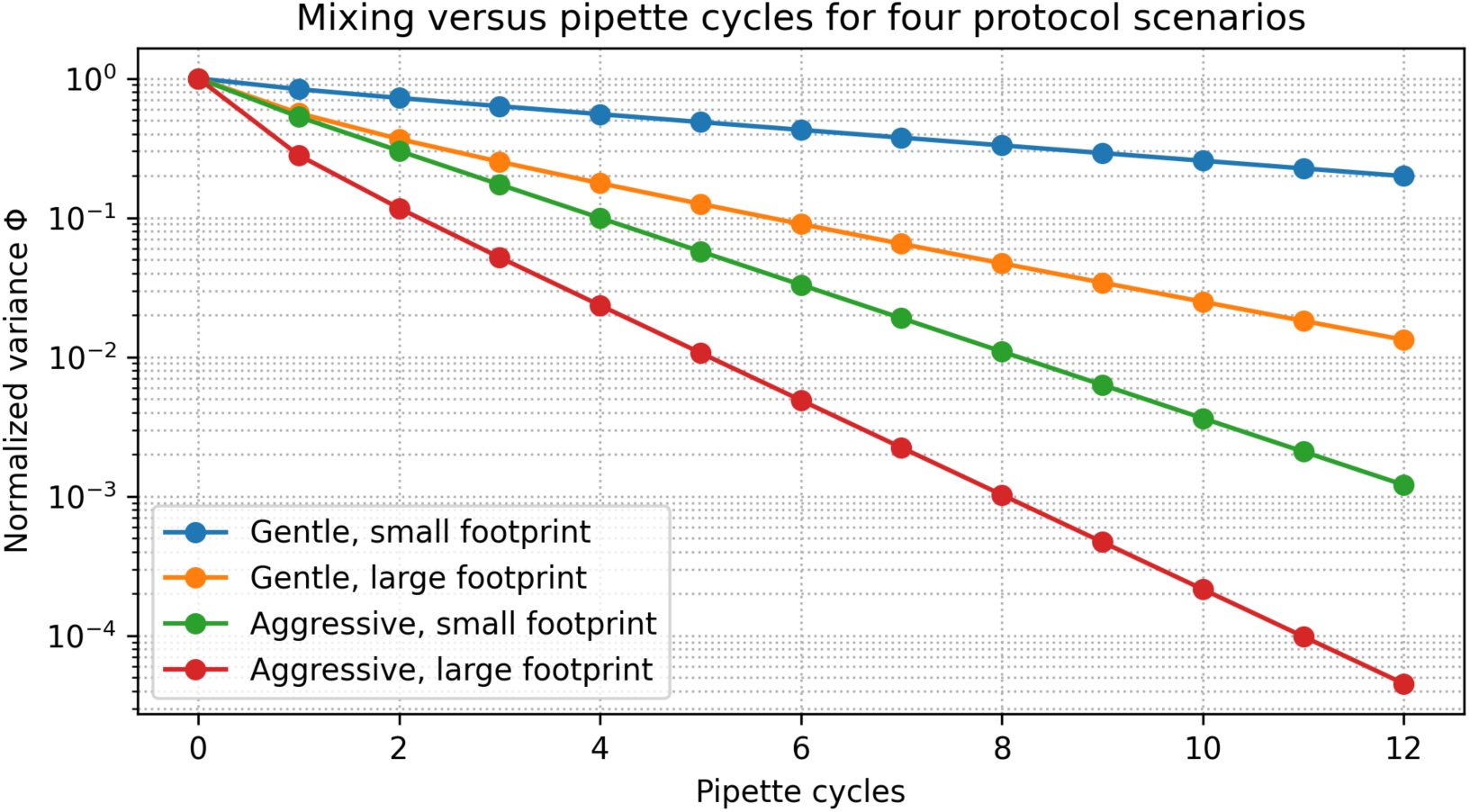
Mixing versus pipette cycles for four protocol scenarios. Normalized concentration variance Φ as a function of pipette cycle number for four parameter regimes in the reduced-order mixing model. Each curve corresponds to a different combination of jet strength and jet footprint size: gentle/small footprint, gentle/large footprint, aggressive/small footprint, and aggressive/large footprint. All simulations start from a biphasic initial condition with the top half of the well at high solute concentration and the bottom half at low concentration. Aggressive, large-footprint jets achieve rapid homogenization (steep decay in Φ), whereas gentle, small-footprint jets mix slowly and may not reach Φ ≤ 0.05 within 10–12 cycles. This hierarchy of mixing efficiency is qualitatively consistent with expectations from microtiter hydrodynamics and CFD studies.

By contrast, gentle jets with small footprints (scenario 1) often remain poorly mixed even after 10–12 cycles, with Φ remaining at values well above 0.05. Increasing the footprint (scenario 2) substantially improves performance, but gentle, large-footprint jets still generally require more cycles than their aggressive counterparts to reach Φ ≤ 0.05.

This hierarchy—fast mixing for high-power, broadly engaging agitation vs slow or incomplete mixing for low-power, localized agitation—is consistent with trends seen in shaken microwell systems and small-scale bioreactors [3,4,6,8,9,12,13]. It supports the expectation that pipette mixing efficiency depends not only on stroke count but also on the effective “power input” and volume fraction sampled per stroke.

### Effect of pipette angle and tip placement

Next, the influence of pipette angle and tip placement on mixing rate was examined. **Figure 3** shows Φ versus cycle number for a representative aggressive-jet scenario at three angles: 0° (vertical), 30°, and 60° from vertical, with the tip located at the well center in all cases. Across this moderate angle range (0–60°), the model predicts only modest differences—typically one cycle or less in the number required to reach Φ ≤ 0.05. Tilted configurations tend to perform as well as or slightly better than the purely vertical one, likely because the jet footprint sweeps a larger horizontal area.

**Figure 3.**
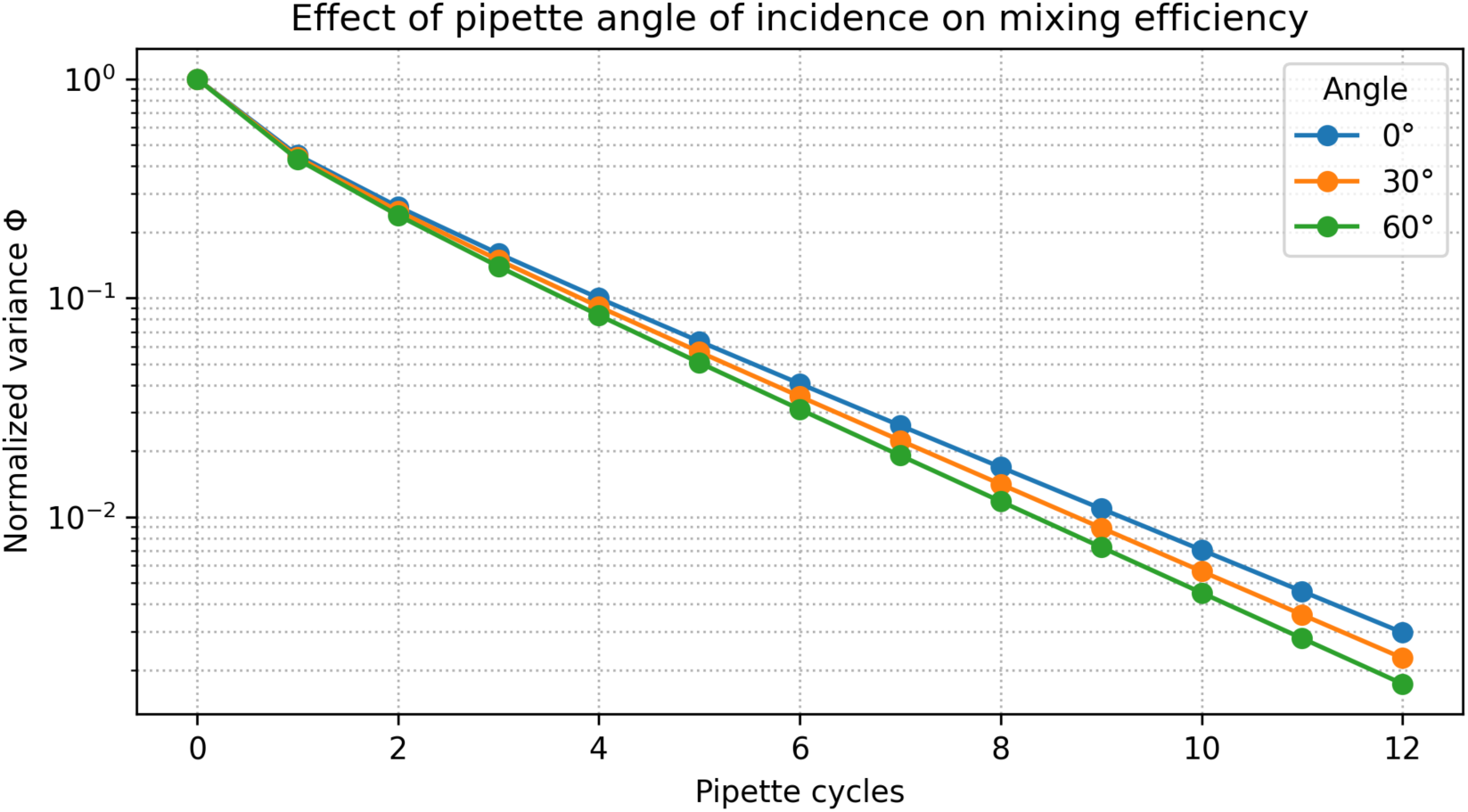
Effect of pipette angle of incidence on mixing efficiency. Normalized concentration variance Φ versus pipette cycle number for an aggressive jet configuration at three pipette angles relative to vertical (0°, 30°, and 60°). The pipette tip remains centered in the well in all cases, and only the angle is varied. Across this range, the model predicts only modest differences—typically at most one cycle—in the number required to reach Φ ≤ 0.05. Moderate tilting (30° or 60°) produces small improvements in later mixing compared to the purely vertical configuration, but all angles show similar Φ values over the first few cycles, indicating that pipette angle is a second-order factor compared with jet strength and footprint size.

**Figure 4** compares centered versus off-center tip placement for a fixed angle and jet strength. In this reduced-order model, modest off-center pipetting generally yields a slightly faster decay in Φ than the centered configuration (typically about one fewer cycle to reach Φ≤0.05), while both configurations eventually achieve similar degrees of mixing. These results qualitatively mirror observations in CFD studies where the location and extent of the main energy input influence how quickly stagnant regions are eroded, although the present model does not resolve detailed recirculation zones [4,7,9,12].

**Figure 4.**
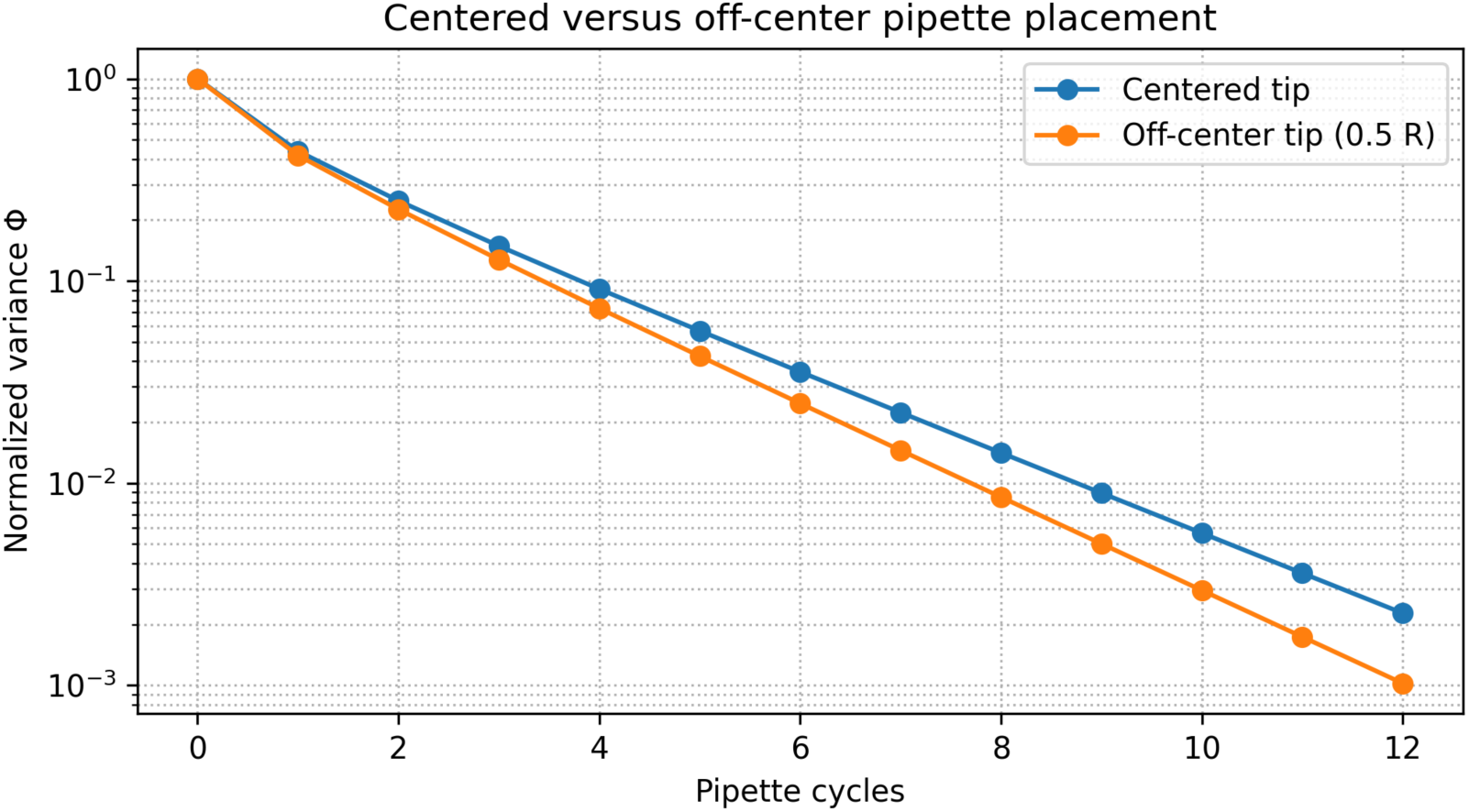
Centered versus off-center pipette placement. Comparison of mixing dynamics for a fixed jet strength, footprint, and angle, with the pipette tip placed either at the center of the well (centered) or offset toward the wall (off-center). Curves show normalized concentration variance Φ versus pipette cycle number. In this reduced-order model, modest off-center placement yields a slightly faster decay in Φ (typically about one fewer cycle to reach Φ≤0.05), although the differences are modest and both configurations ultimately achieve similar levels of homogeneity. These results highlight that tip placement can influence mixing efficiency, but that its impact is smaller than that of jet strength and footprint size.

Together, these simulations suggest that pipette angle within typical manual ranges is a second-order factor compared to jet strength and footprint, and that engaging a large fraction of the well volume is more important than strict centering. In the present model, moderate off-center placement performs as well as, or slightly better than, a centered tip, whereas strongly localized mixing near a single corner (not explicitly resolved here) would be expected from CFD results to leave more persistent stagnant regions.

**Figure 5** and **Table 1** summarize a small parameter sweep exploring how jet strength and jet footprint radius jointly control the number of cycles required to reach a practical mixing threshold (Φ ≤ 0.05). In this reduced-order model, increasing either the jet strength (vertical axis) or the footprint radius (horizontal axis) systematically shifts the system toward faster mixing, as reflected by lower cycle counts in the heat map. Weak, highly localized jets (lower left) require the largest number of cycles and in some cases do not achieve the threshold by the end of the simulated 12-cycle window, whereas intermediate conditions (e.g., moderate strength with a modest footprint) typically reach Φ ≤ 0.05 in roughly 6–10 cycles. The most aggressive, broadly sweeping jets (upper right) achieve effective mixing in only ∼3–5 cycles. Overall, **Figure 5** makes explicit that the widely used “three-cycle” heuristic is only valid near the high-strength, large-footprint corner of parameter space; for gentler jets or more restricted footprints, substantially more cycles are required to reach comparable levels of homogeneity.

**Figure 5.**
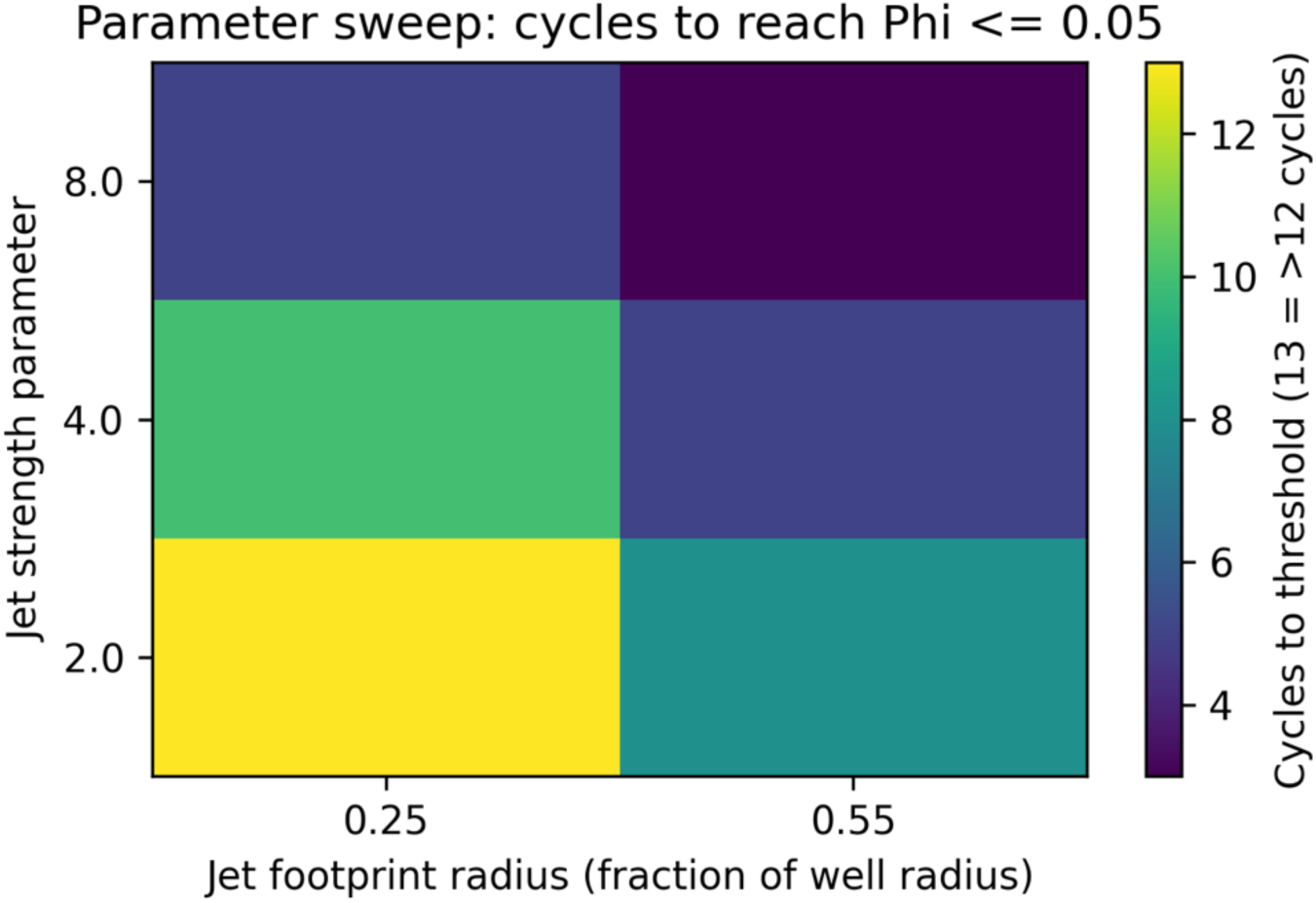
Parameter sweep reveals regimes of slow and fast mixing. Heat map showing the number of pipette cycles required to reach a practical mixing threshold (Φ ≤ 0.05) as a function of jet strength (vertical axis) and jet footprint radius (horizontal axis) in the reduced-order macrocell model. Each cell represents the first cycle at which the normalized concentration variance falls below the threshold; darker colors indicate fewer cycles (faster mixing), and lighter colors indicate more cycles (slower mixing). Aggressive jets with larger footprints (upper right) typically achieve adequate mixing within 3–5 cycles, whereas weaker and more localized jets (lower left) may require more than 10 cycles to reach the same degree of homogeneity.

**Table 1.**
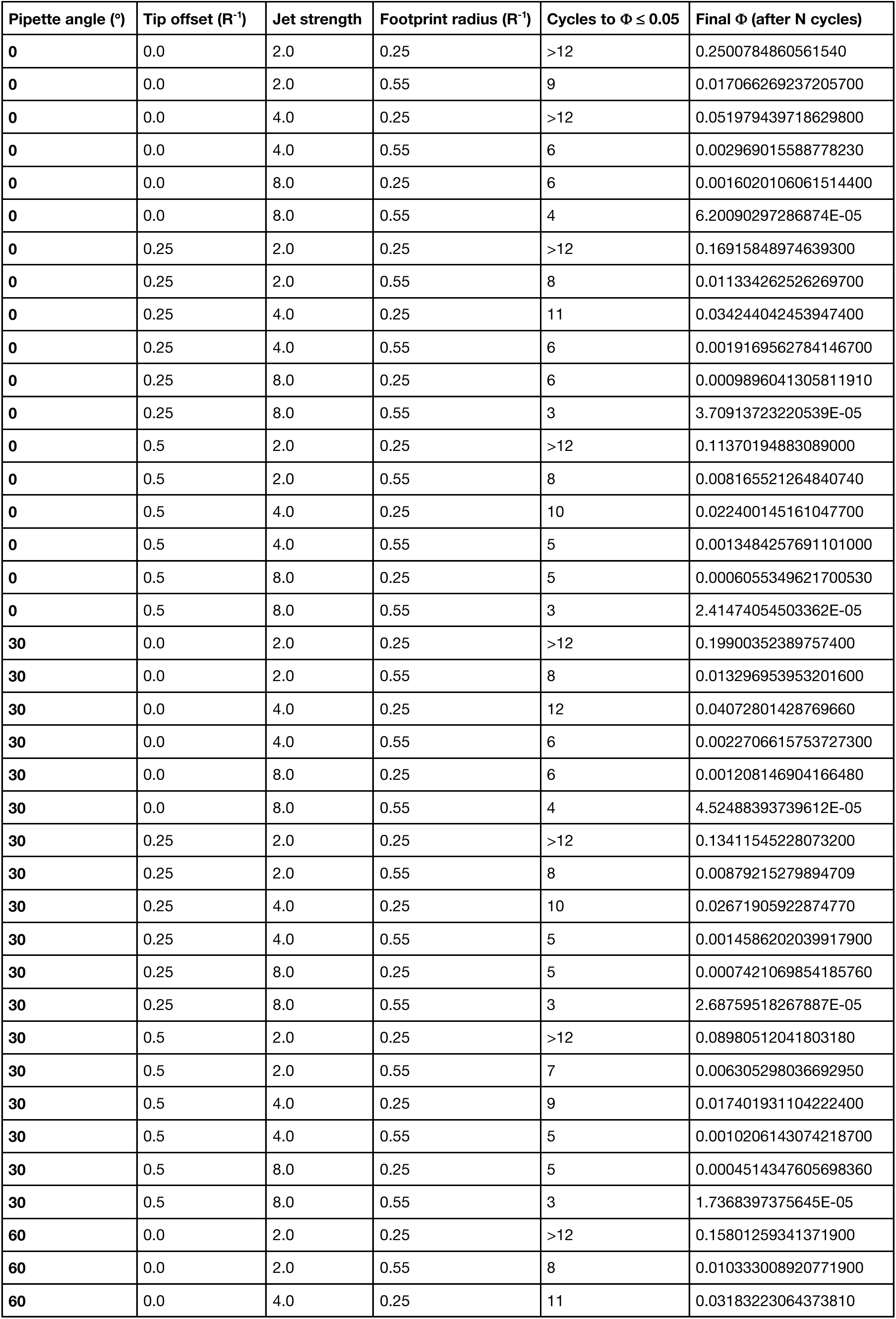
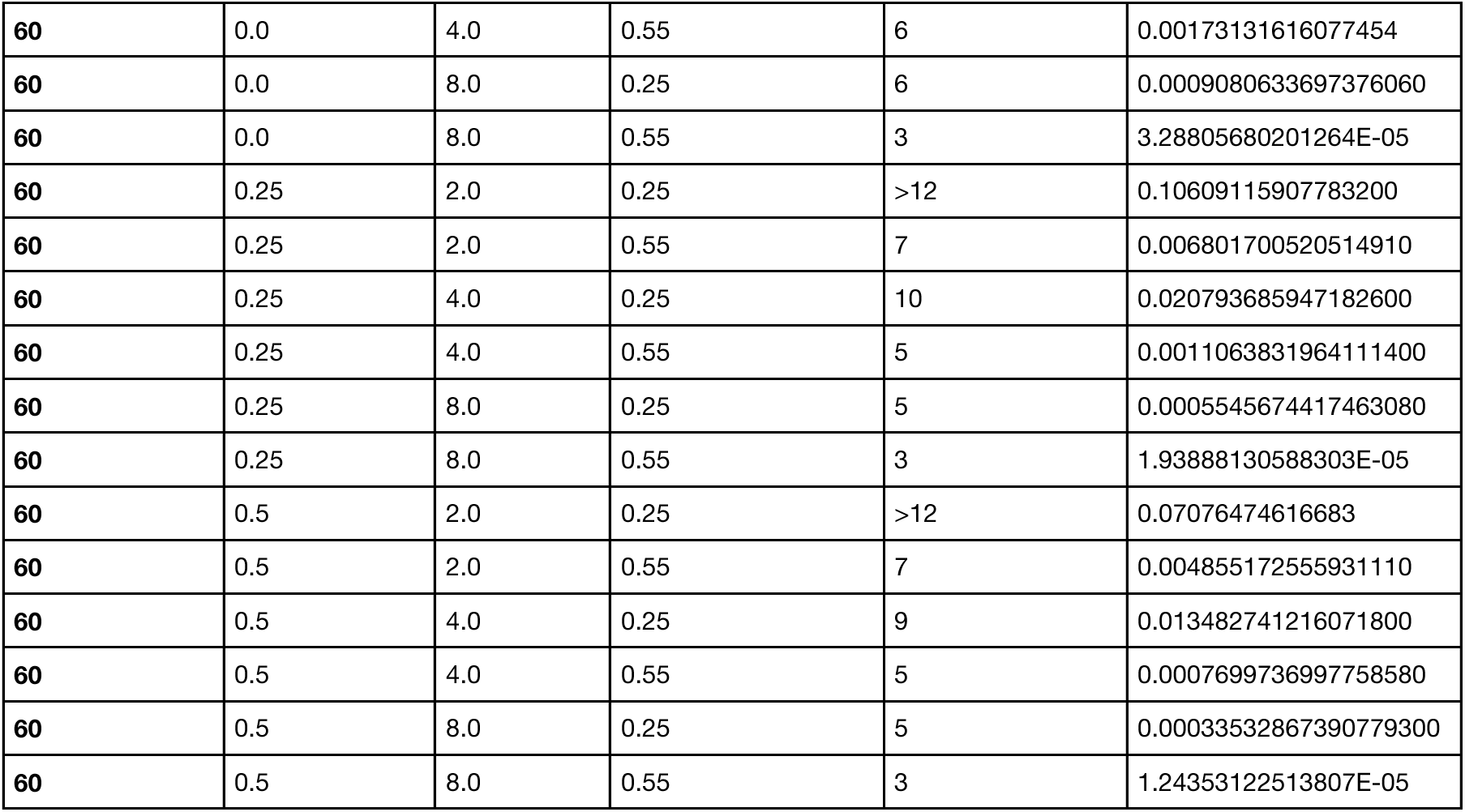
Parameter sweep of pipette-driven mixing in a 96-well macrocell model. Results of simulations varying tip radial offset, jet strength, and jet footprint radius at a fixed pipette angle. For each configuration, the table reports the number of cycles required to reach the mixing threshold (Φ≤0.05) and the final normalized variance Φ after 12 cycles when the threshold is not reached earlier.

### Stroke counts required for practical mixing thresholds

**Table 2** summarizes the approximate number of cycles required to achieve Φ ≤ 0.05 (“well mixed”) for representative combinations of jet strength, footprint size, and pipette configuration. Although exact numbers depend on the model parameter values, several robust patterns emerge:

- For aggressive jets with reasonably large footprints, Φ ≤ 0.05 is typically achieved within 3–6 cycles, and Φ ≤ 0.01 within 6–10 cycles.
- For gentle jets with optimal (large) footprints, 6–10 cycles may be needed to reach Φ ≤ 0.05, and achieving Φ ≤ 0.01 within a practical number of cycles (≤20) is not guaranteed.
- For gentle, small-footprint configurations, particularly with off-center tips, even 10–12 cycles may leave significant residual heterogeneity.

**Table 2.**
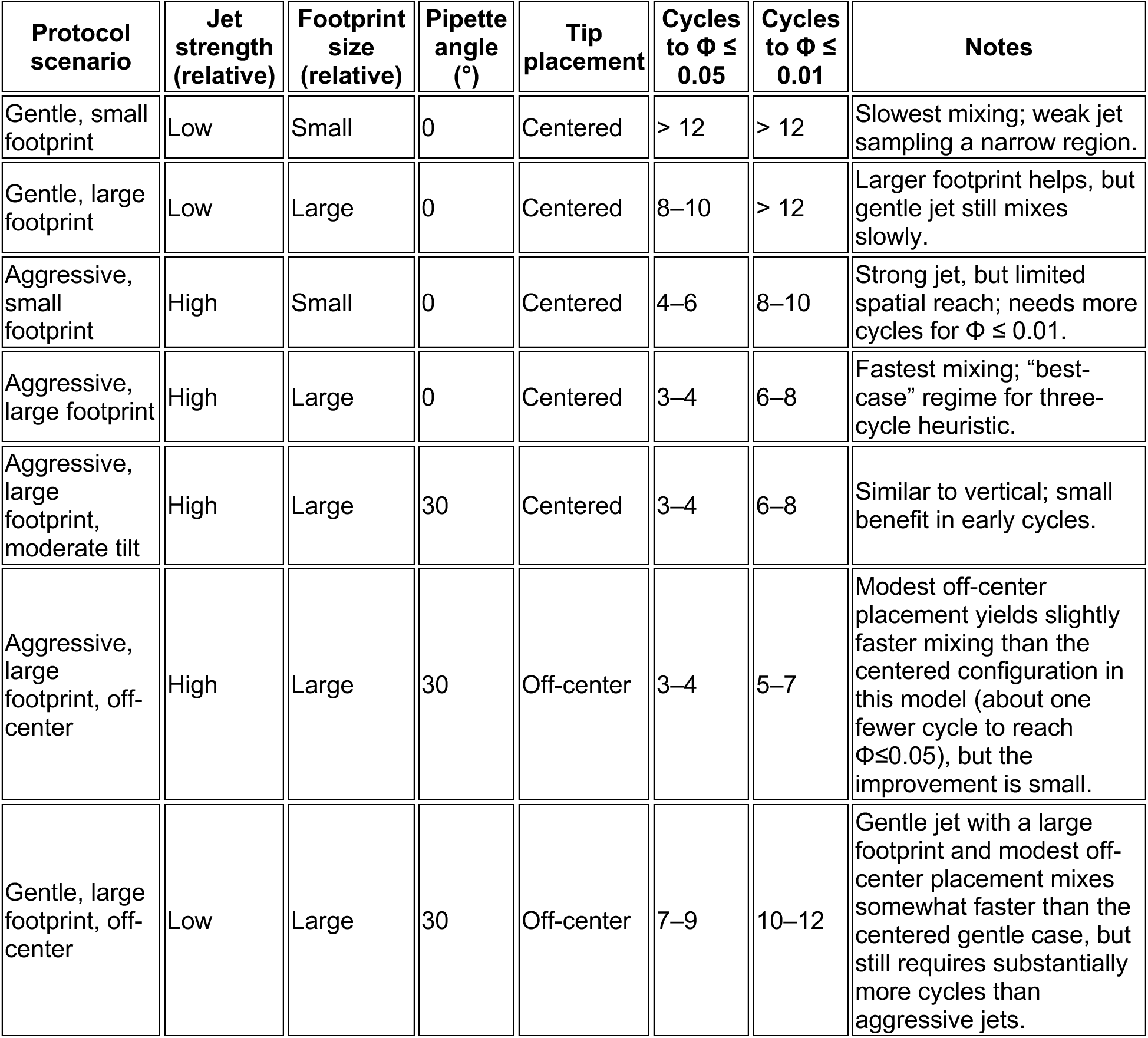
Model-predicted cycles required to reach practical mixing thresholds under representative pipette configurations. Number of pipette cycles required to achieve normalized concentration variance Φ≤0.05 (“well mixed”) and Φ≤0.01 (“highly mixed”) for selected combinations of jet strength, jet footprint size, pipette angle, and tip placement in the reduced-order model. Rows correspond to representative protocol scenarios (e.g., gentle/strong jet, small/large footprint; centered vs off-center tip), and columns report the cycles needed to reach each threshold or “>12” if the threshold is not reached within the simulated 12-cycle window. The underlying parameter sweep and set of combinations are provided in **Table 1**.

Thus, the commonly cited rule of thumb that “three cycles are enough” is not consistently valid. It may hold in favorable situations where a relatively powerful jet is used, the pipette volume fraction is large, and the footprint samples most of the well volume. But in gentler, more localized scenarios—which may arise, for example, when trying to avoid bubble formation or cell damage—significantly more cycles may be required.

Because this model is linear and fast to evaluate, investigators can in principle tune parameters to match their specific pipette settings (e.g., stroke volume, speed) and plate geometries to obtain more quantitative predictions of required stroke counts. For the purposes of this study, qualitative trends and order-of-magnitude assessments remained the focus.

### Comparison to hydrodynamics and CFD studies

The current reduced-order model simplifies the detailed velocity fields, yet its behavior aligns with several key observations from more detailed hydrodynamics and CFD investigations:

- <u>Power input and mixing time:</u> Dürauer et al. and Montes-Serrano et al. showed that specific power input and energy dissipation correlate inversely with mixing times in shaken microtiter plates [4,6]. In the current model, increasing jet strength 𝐾_jet_ and footprint size plays an analogous role, reducing the number of cycles required to decrease Φ below a given threshold.
- <u>Geometric sensitivity:</u> Engineering characterization of shaken 24- and 96-well plates by Zhang et al. highlighted substantial differences in flow regimes and mixing behavior between geometries [9]. The current parameter sweeps over footprint size and tip placement similarly show that subtle geometric changes can have quantifiable effects on mixing efficiency.
- <u>Consequences of inadequate mixing:</u> In permeability studies using 96-well sandwich plates, Eriksen et al. found that orbital shaking sometimes failed to provide adequate mixing, leading to biased permeabilities, whereas magnetic stirring was more effective [14]. The present results reinforce the message that insufficient agitation (in the present case, weak pipette jets and too few strokes) can leave residual gradients that would bias assay readouts, particularly for diffusion-limited processes.

Given the relatively coarse resolution of the macrocell grid and the simplified jet representation, one may stop short of claiming quantitative predictive accuracy. However, the model’s qualitative behaviors are consistent with accepted principles of microtiter hydrodynamics and previously reported mixing hierarchies [3,4,6–9,12–14], suggesting that it can serve as a practical design tool for pipette mixing protocols.

### Online survey reveals a wide range of stroke counts

The survey of online sources yielded 42 protocols or posts with explicit pipette-mixing stroke counts in 96-well plates or 96-well PCR plates, drawn from the Reddit.com/LabRats subreddit, commercial kit manuals, and a few peer-reviewed articles (**Table 3**). When a range of stroke counts was included in the protocol or online forum post, the midpoint was recorded for the purposes of data display. The distribution of stroke counts is shown in **Figure 6**.

**Figure 6.**
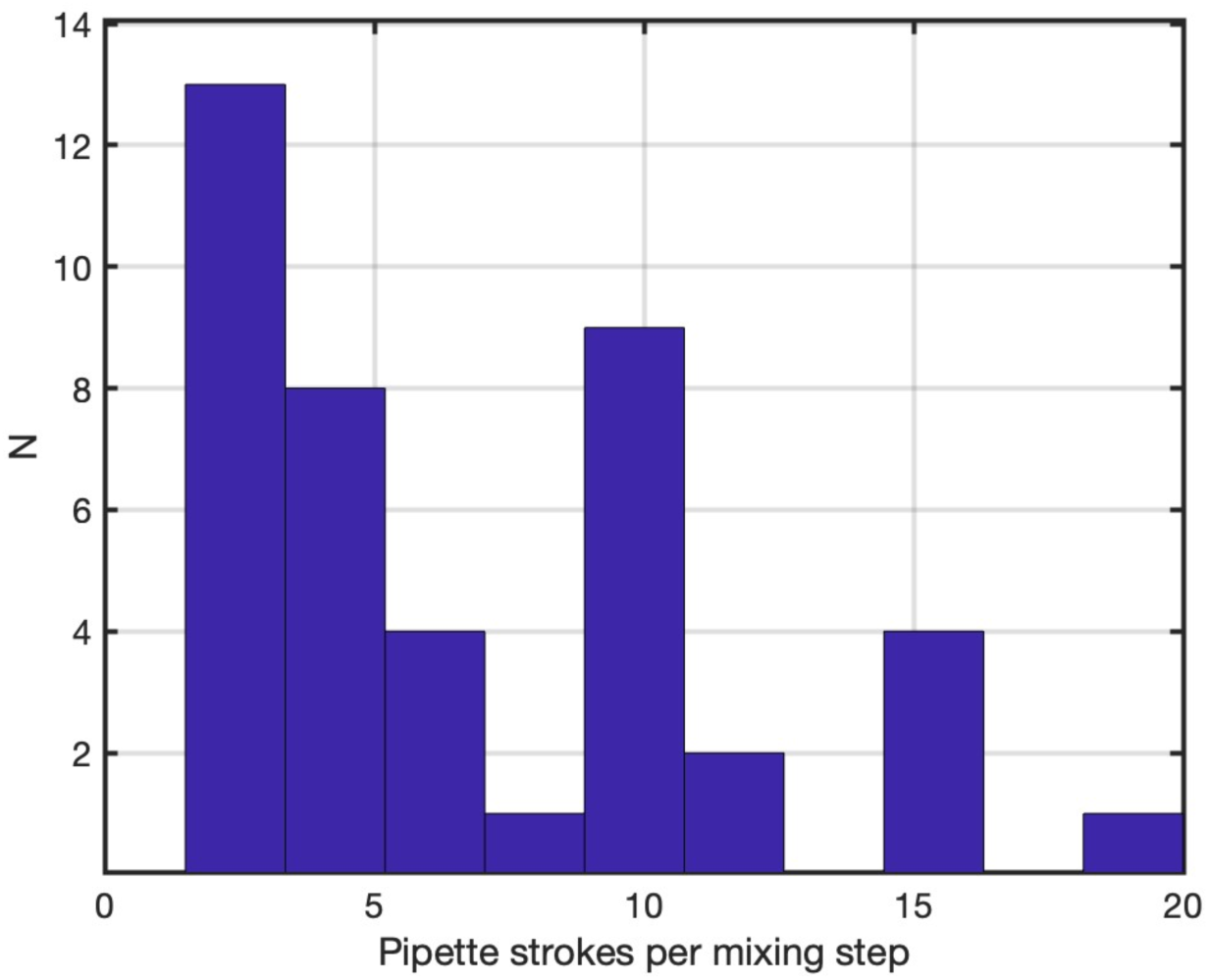
Distribution of pipette mixing stroke counts used in 96-well protocols. Histogram of the number of “pipetted up and down *N* times” strokes per mixing step extracted from 96-well protocols in online research forums, vendor documentation, and published articles (42 protocol instances in total). The dataset (**Table 3**) includes those sources that explicitly specify a stroke count. Many protocols do not report stroke counts and are therefore not represented in the graph.

**Table 3.** Pipette mixing in 96-well plate protocols found via internet search. While most published protocols involving mixing in well plates by pipette withdrawal and ejection cycles do not specify the number of mixing cycles, the data presented here were found on the Reddit.com “LabRats” subreddit, in vendor assay kit documentation, and 3 peer-reviewed protocol papers. Note that in a limited number (n=4) cases, the protocol also specified a pipetting volume, expressed as a percentage of the total liquid volume contained in the well.

| Protocol source | Pipetting stroke count | Pipette volume |
| --- | --- | --- |
| https://www.reddit.com/r/labrats/comments/1oezr1p/do_you_mix_by_pipetting_in_and_out_after_adding/ | "couple of times" (=2) | 80% |
|  | 2 |  |
|  | 10 |  |
|  | 20 |  |
| https://www.reddit.com/r/labrats/comments/1kyfnn0/how_do_you_guys_mix_your_qpcr_plates/ | 10 |  |
|  | 1 – 2 |  |
|  | 6 | 80% |
|  | 1 – 2 |  |
| https://www.reddit.com/r/labrats/comments/f20obn/how_do_you_guys_mix_your_96_well_plates/ | 2 – 3 |  |
|  | 2 – 3 |  |
| https://www.reddit.com/r/labrats/comments/1g1qkpt/tips_for_equal_seeding_in_a_96well_plate/ | 2 – 3 |  |
| https://www.reddit.com/r/labrats/comments/1l5kpg3/any_tips_of_removing_air_bubbles_forming_in_the/ |  | 25-50% |
|  | 10, 15 | 50% |
| https://www.anamed.com.tr/wp-content/uploads/2022/11/Reproducibility_Serial_Dilutions_V01.pdf | 5 |  |
| https://www.cambio.co.uk/library/images/html_images/Aline/Protocol-PureGenome-Blood-gDNA-v03.pdf | 10 |  |
| https://norgenbiotech.com/sites/default/files/resources/PI62400_6_Plant_DNA_Isolation_96_Well_Kit_Magnetic_Bead_System_Insert.pdf?srsltid=AfmBOooM3J10ZXTltbZpZroHwTzADd1H8fuUV7g7Djlib-iPlb8b-4Qj | 10 |  |
| https://www.researchgate.net/post/How-should-I-pipette-2ul-of-cDNA-without-making-bubbles | 2 |  |
|  | 5 – 10 |  |
| https://nanoporetech.com/document/ligation-sequencing-gdna-pcr-barcoding-sqk-lsk114-with-exp | 10 – 20 |  |
| https://www.protocols.io/view/protoplast-isolation-and-transfection-in-a-96-well-ccc4ssyw.pdf | 5 |  |
| https://documents.thermofisher.com/TFS-Assets/LSG/manuals/sequalprep_platekit_man.pdf | 5 |  |
| https://www.integra-biosciences.com/sites/default/files/documents/498-23_Website_Application_Guide_Serial_Dilution_150623.pdf | 3 |  |
| https://content.protocols.io/files/paffb8y37.pdf | 10 – 20 |  |
| https://www.integra-biosciences.com/united-states/en/applications/fast-and-efficient-automated-sample-transfer-tubes-plates-assist-plus-pipetting-robot | 3 |  |
| https://www.integra-biosciences.com/global/en/blog/article/how-do-serial-dilutions-including-calculations | 5 |  |
| https://www.thermofisher.com/us/en/home/references/protocols/cell-and-tissue-analysis/elisa-protocol/elisa-sample-preparation-protocols/sample-preparation-adherent-suspension-cells.html | 5 – 6 |  |
| https://www.magbiogenomics.com/pub/media/productattach/h/i/highprep_plasmid_dna_-_protocol.pdf?srsltid=AfmBOorzdsUi2iriepRTjxeYJk00psmjMWiHcnwzeVu1cDbjYJPafzN | 10 |  |
| https://www.pacb.com/wp-content/uploads/Procedure-checklist-Generating-Pure-Target-libraries-with-Pure-Target-kit-96-automation-protocol.pdf | 5 |  |
| https://www.agilent.com/cs/library/usermanuals/public/5991-7930EN.pdf?srsltid=AfmBOoot1e9GIUafwI3sD9TzTV5OHBuwD919Lz38spZ5uwBj4yS5gXCi | 3 – 5 |  |
| https://www.beckman.com/reagents/genomic/cleanup-and-size-selection/pcr/ampure-xp-protocol | 10 |  |
| https://www.sopachem.com/lifesciences/wp-content/uploads/2018/10/Instruction-Guide-for-use-of-Syntrix-P.pdf | 10 – 20 |  |
| https://genome.med.harvard.edu/documents/sequencing/Agencourt_CosMCPrepProtocol.pdf | 10 |  |
| https://www.neb.com/en-us/protocols/2018/06/04/protocol-for-nebnext-direct-custom-ready-panels-neb-e6631?srsltid=AfmBOoorf52ORMX9w-CzwxdW44lt0xp6_1AczkCw2QRyQ5scAUE-ZQH2 | 10 |  |
| https://www.bdbiosciences.com/en-eu/resources/protocols/96-deep-well-staining-cytokine | 3 |  |
| https://nanoporetech.com/zh/document/ligation-sequencing-gdna-pcr-barcoding-sqk-lsk114-with-exp | 10 – 20 |  |
| https://www.protocols.io/view/protoplast-isolation-and-transfection-in-a-96-well-yxmvnm1d6g3p/v1 | 5 |  |
| https://protocols.opentrons.com/categories/Sample%20Prep/Serial%20Dilution | 12 |  |
| https://www.andrewalliance.com/wp-content/uploads/2022/02/618.002_IQ-OQ-Pipette-v7-NV-20220225.pdf | 2 |  |
| https://www.sciencedirect.com/science/article/pii/S2472630322020659 | 3 |  |
| https://documents.thermofisher.com/TFS-Assets/LSG/manuals/100008679.pdf | 6 |  |
| https://www.jove.com/v/64298/fast-colony-forming-unit-counting-96-well-plate-format-applied-to | 10 |  |
| https://www.jove.com/t/57713/diagonal-method-to-measure-synergy-among-any-number-of-drugs | 5 |  |
| https://www.nature.com/articles/s41598-018-30305-z | 6 |  |

It is noted that the recommended stroke count varies broadly from 2 – 20, with distinct peaks found at 2.5 (corresponding to a specified range of 2 – 3), another peak at 10, and a third peak at 15 (corresponding to a specified range of 10 – 20). The 10-stroke specification is particularly common across diverse assay types, including cell-based assays, bead or pellet resuspension, enzyme assays, and PCR or library-preparation steps. A smaller subset of protocols, especially vendor kits and some other researcher recommendations, prescribe ∼20 strokes for more demanding mixing steps (e.g., dense pellets, viscous suspensions, or critical serial dilutions).

Many otherwise detailed protocols still use qualitative language (“pipette several times to mix,” “mix thoroughly”) without specifying stroke counts, so this dataset greatly underestimates the total number of assays where pipette mixing plays an important role. The entries captured here therefore represent a conservative sample of cases where researchers judged stroke count important enough to report explicitly.

When interpreted through the lens of the reduced-order model, this distribution is reasonable. Stroke counts of 10–12 align well with regimes where Φ ≤ 0.05 can be achieved under moderately aggressive, well-placed jets, while 3–5 strokes are adequate primarily for simpler mixing tasks or more powerful jets. Stroke counts of 20+ offer a substantial safety margin for difficult suspensions. The survey thus supports the main conclusion: the informal “three strokes are enough” expectation reflects only one end of a much broader, empirically used spectrum of pipette-mixing efforts.

### Recommendations for 96-well pipette mixing

- <u>Do not assume three strokes are always enough.</u> The present model suggests that three cycles only suffice in favorable regimes (relatively strong jets and broad footprints). For many realistic scenarios, especially gentle pipetting or viscous suspensions, more strokes will be required.
- <u>Aim for 5–10 strokes as a default for challenging mixes.</u> This range is both common in the literature and consistent with model-predicted stroke counts needed to reach Φ ≤ 0.05 under moderately aggressive conditions.
- <u>Use more strokes (up to ∼20) for difficult suspensions.</u> For bead-rich, highly concentrated, or viscous samples, 10–20 strokes are often used in published protocols and are consistent with the current simulation as a way to guard against residual gradients.
- <u>Center the pipette tip or use only moderate offsets.</u> In the present model, modest off-center placements perform as well as, or slightly better than, a centered tip, but in practice centering is a simple default that helps distribute mixing energy symmetrically and avoids extreme localization near one corner of the well.
- <u>Use pipette angle as a fine-tuning parameter, not a primary knob.</u> Within typical manual ranges (0–60°), angle has a smaller effect than jet strength and footprint, but gentle tilting can modestly improve mixing.
- <u>Standardize and report stroke counts.</u> Given the impact of mixing on assay readouts, investigators should treat stroke count as a controlled parameter and report it explicitly, particularly in high-throughput or diffusion-limited assays.

These recommendations are deliberately conservative, reflecting both the model’s predictions and the clustered stroke counts observed in the literature. They are intended as a starting point rather than a one-size-fits-all prescription.

### GPT-native modeling and responsible use of LLMs

A distinctive feature of this work is its GPT-native modeling workflow. A large language model (LLM) was used to help scaffold the Python implementation of the macrocell mixing model, suggest parameterizations, and propose initial text and figures. All code, equations, and interpretations were reviewed, edited, and validated by the human author, who bears full responsibility for the final content.

This mode of collaboration aligns with recent guidance on using ChatGPT and related tools in computational biology and bioinformatics [16–19]. Lubiana et al. outlined practical “tips” for harnessing ChatGPT while emphasizing the need for human oversight and verification [16]. Xu and Wang et al. surveyed emerging applications of LLMs in bioinformatics and biomedical informatics, highlighting both opportunities and limitations [17,18]. Helmy et al. further articulated principles for the careful use of generative AI in scientific workflows [19]. This study contributes a concrete example where an LLM accelerated exploratory modeling and protocol analysis, while the underlying scientific reasoning, model design, and validation remain firmly in the hands of a domain expert.

## Conclusions

A reduced-order fluid model of pipette-driven mixing in 96-well plates was developed and compared to an online search of published pipette-mixing protocols. The model discretizes the well into macrocells with neighbor exchange and represents the pipette jet as a localized enhancement of exchange within a jet footprint aligned with the pipette axis. Despite its simplicity, the model qualitatively reproduces expected mixing hierarchies: aggressive, broad-footprint jets rapidly homogenize the well within a few cycles, whereas gentle, small-footprint jets can require many more cycles and may fail to reach tight mixing thresholds.

Parameter sweeps over jet strength, footprint size, pipette angle, and tip placement revealed that stroke count alone is an incomplete descriptor of mixing quality. The widely used rule of thumb that “three cycles are enough” is only reliable in favorable regimes; in many realistic scenarios, especially gentle pipetting or resuspension of challenging suspensions, more strokes (5–10 or even up to 20) are required to reduce concentration variance to practically negligible levels. Broad jet footprints are consistently beneficial, and tip placement has a modest influence: in the present model, moderate off-center positions perform as well as or slightly better than a strictly centered tip, whereas moderate changes in pipette angle have relatively modest impact.

The online protocol survey, spanning online lab forum posts, vendor kit manuals, and peer-reviewed protocols, shows that published 96-well protocols with explicit pipette-mixing stroke counts cluster around 2–3, and 10 strokes, with a minority of “heavy mixing” steps at 15–20+ strokes. When interpreted through the lens of the present model, these clusters are consistent: 10–12 strokes are a reasonable default for many assays, 20+ strokes provide a safety margin for difficult mixtures, and 3–5 strokes suffice mainly in easy-to-mix or high-power jet scenarios. It is therefore recommended that investigators treat pipette stroke count and configuration as design parameters rather than unexamined habits, and that stroke counts be reported explicitly in methods sections.

Finally, this work illustrates a GPT-native approach to computational biology: a large language model was used as an assistive tool for model scaffolding, code generation, and preliminary text generation, with all outputs curated and validated by the author. As LLMs become more integrated into scientific practice, transparent disclosure and responsible use will be essential. In this case, the synergy between a simple, interpretable physical model, an online collection of real-world practice, and LLM-assisted development enables a straightforward yet rigorous re-examination of a ubiquitous laboratory operation—“pipette, mix, repeat”—and provides actionable guidance for improving the reliability of 96-well plate experiments.

## Acknowledgements

This work was funded in part by National Institutes of Health, grant number R01CA256054 (M.R.K.).

## Appendix: Mixing model implemented in Python

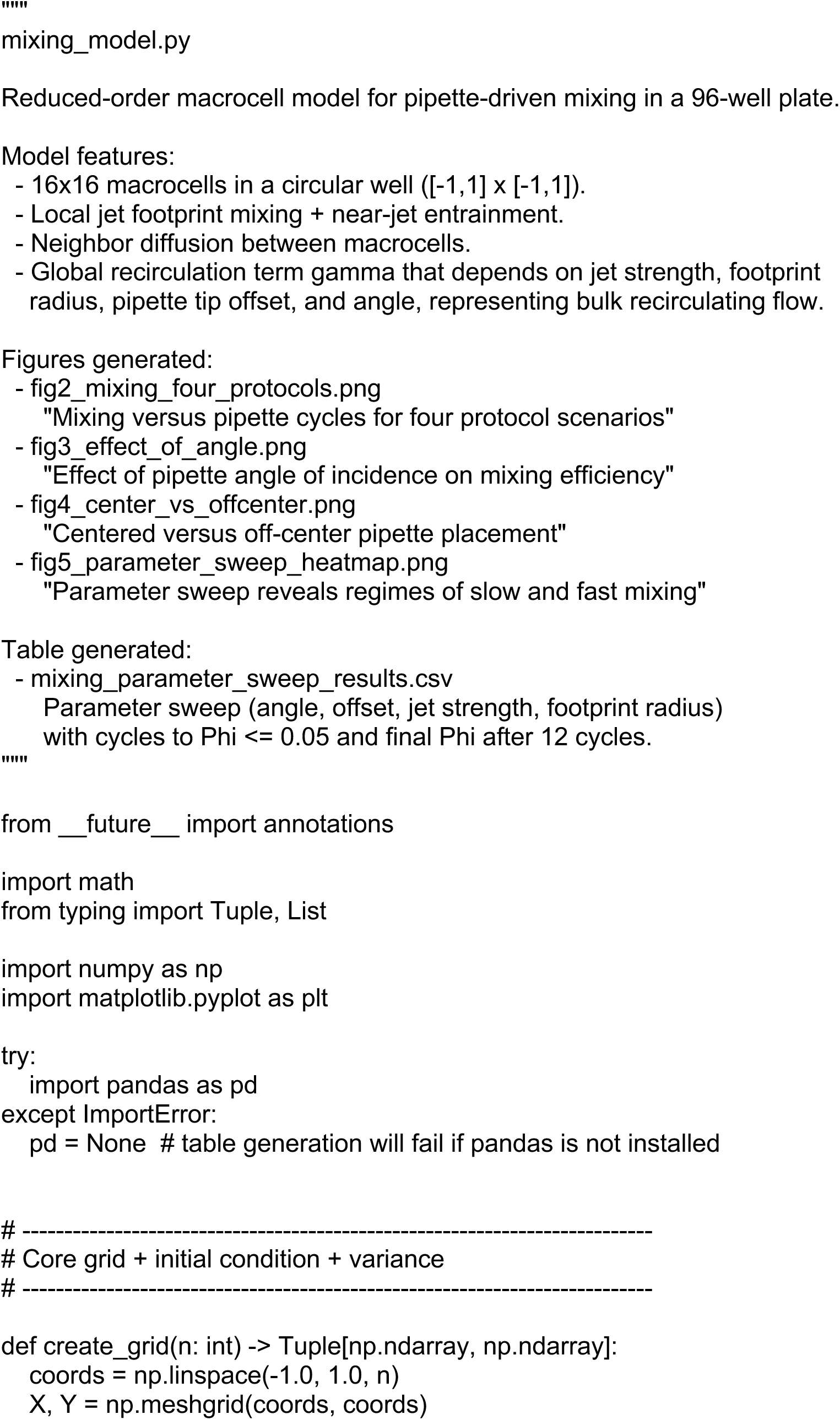
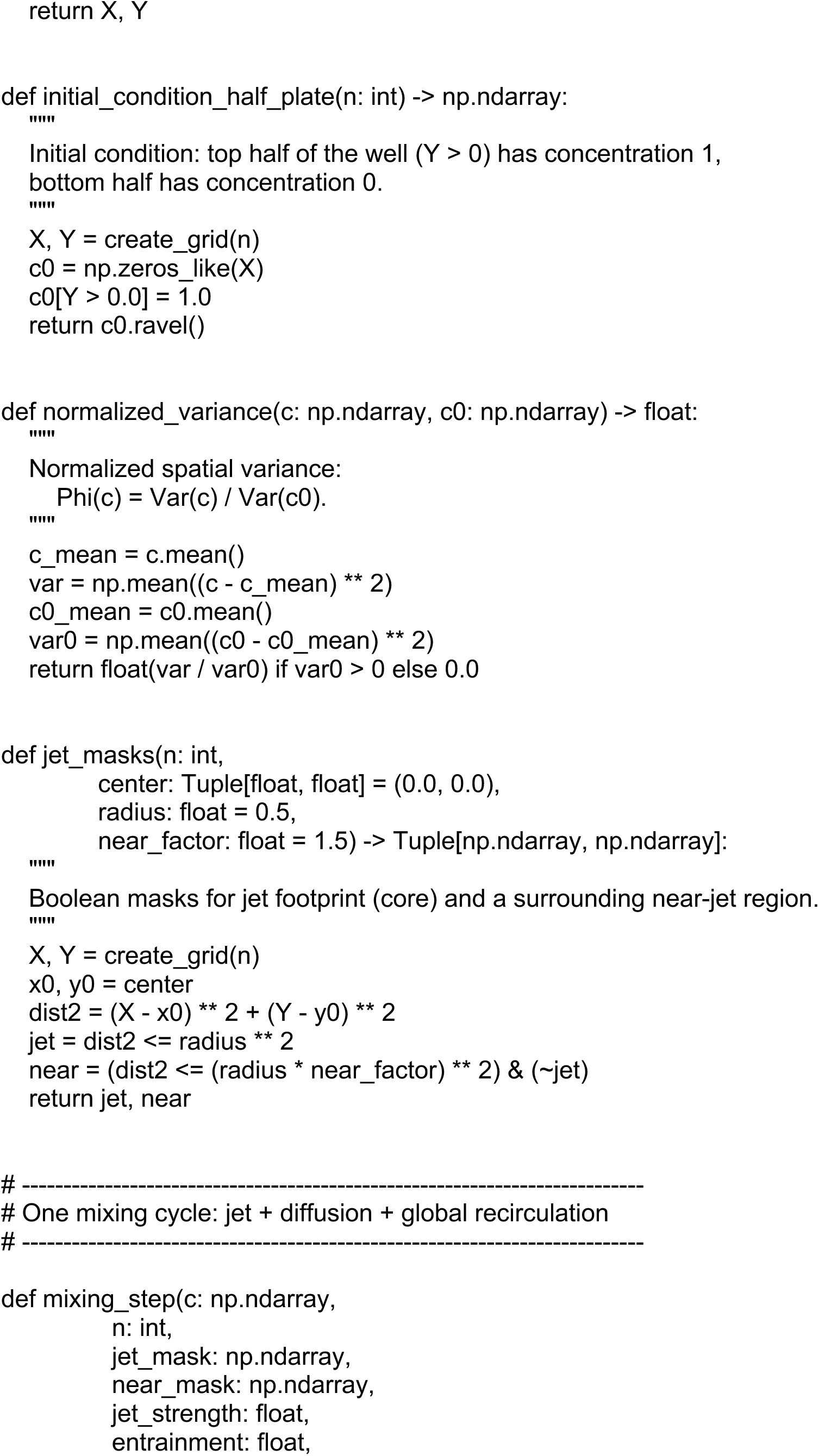
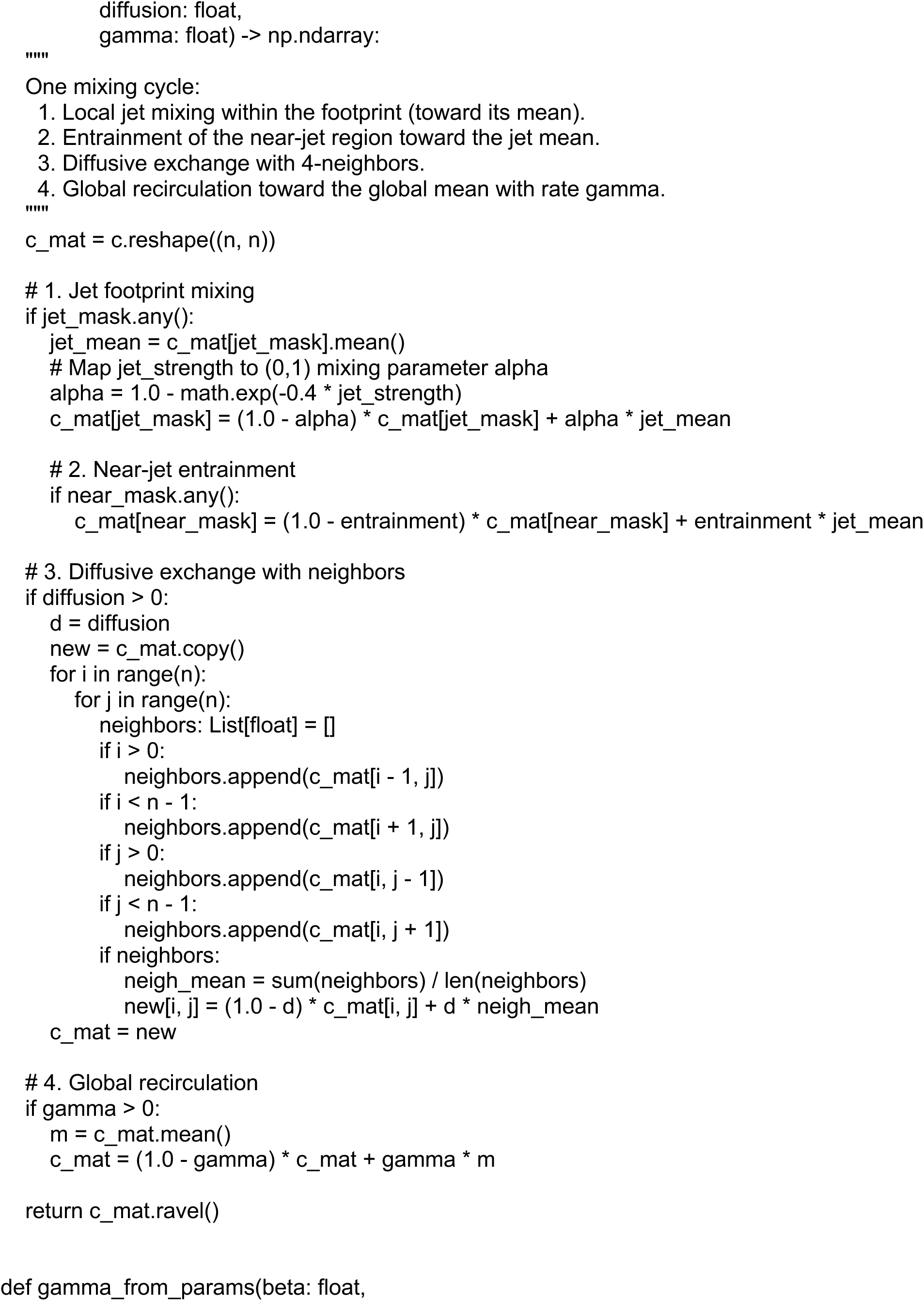
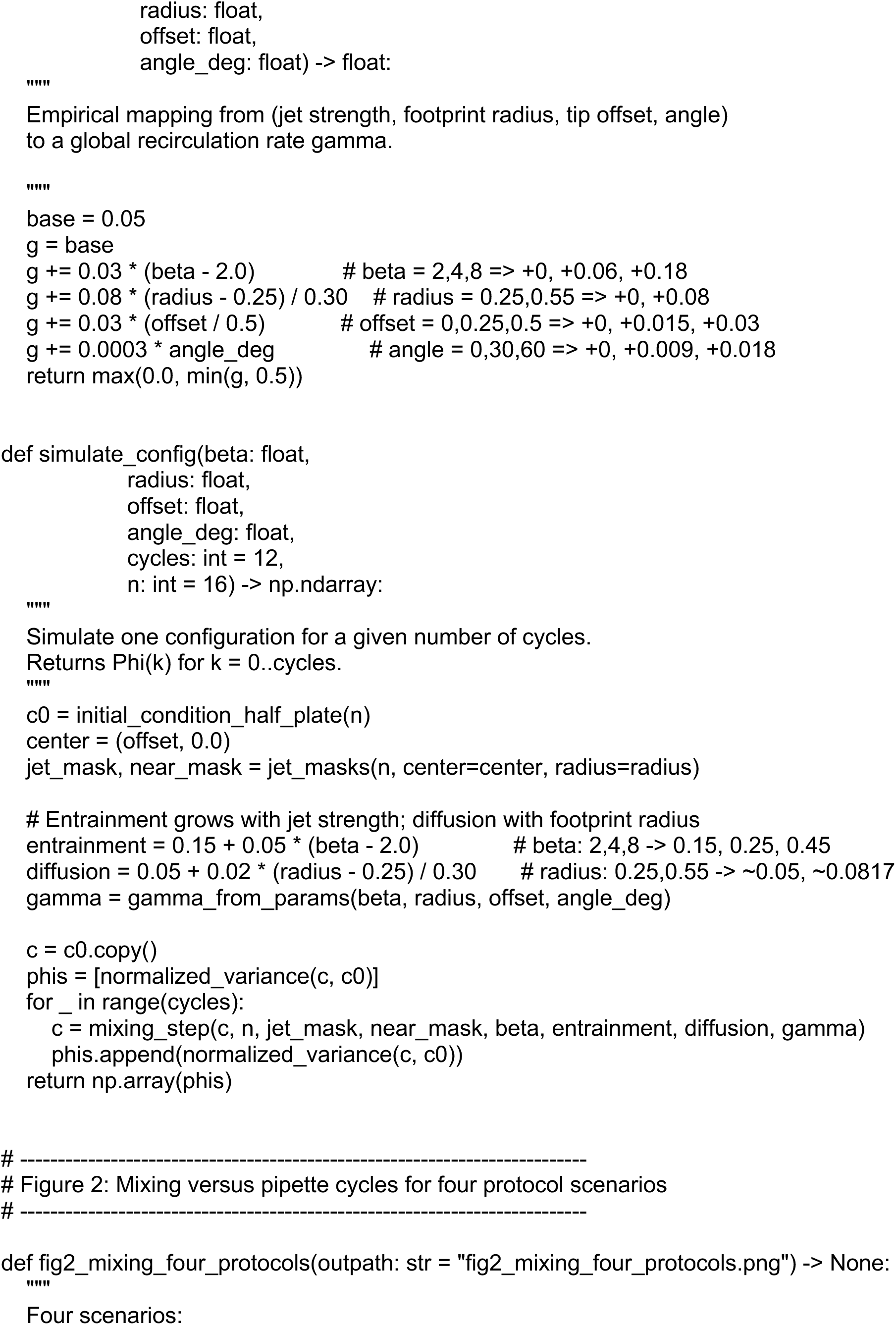
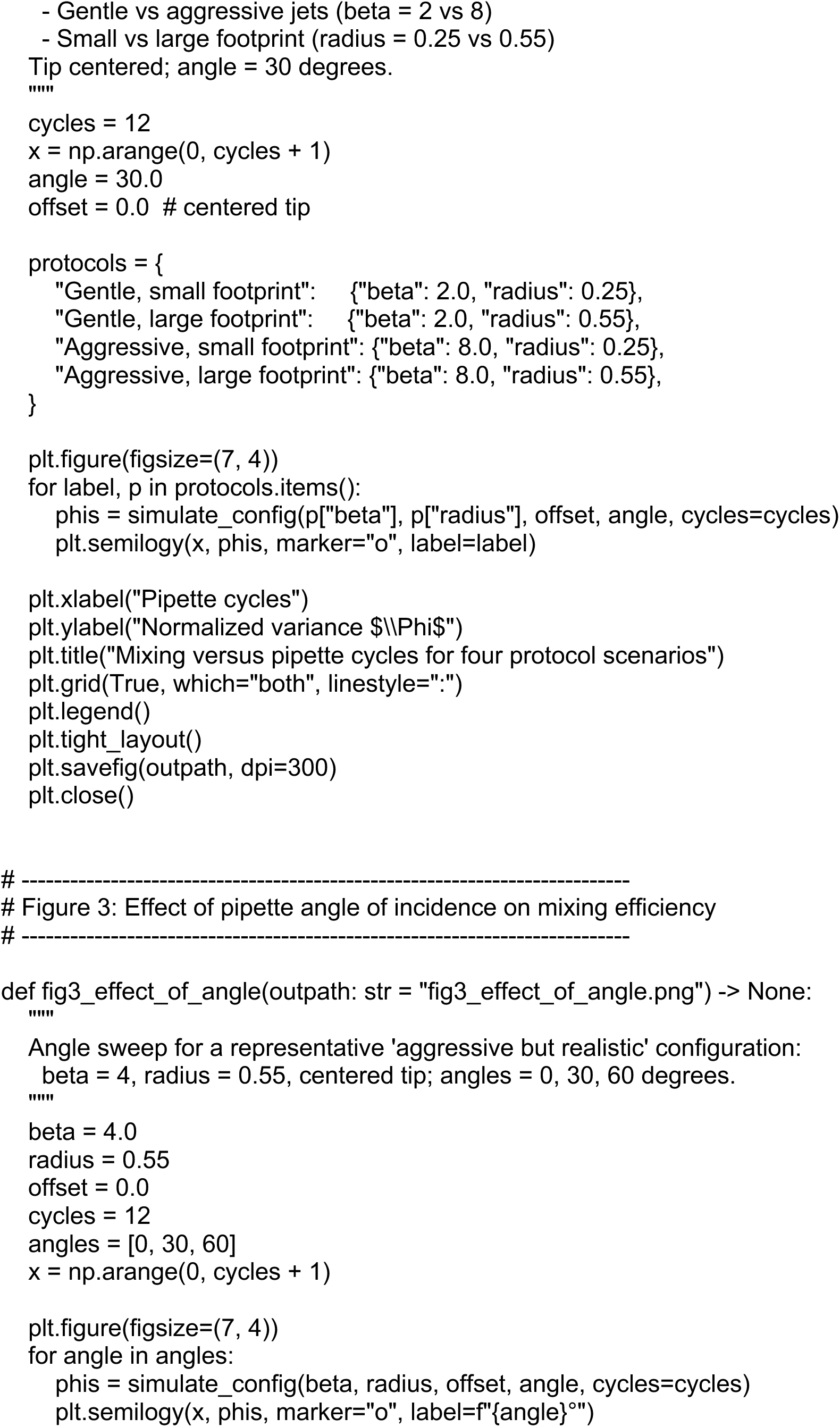
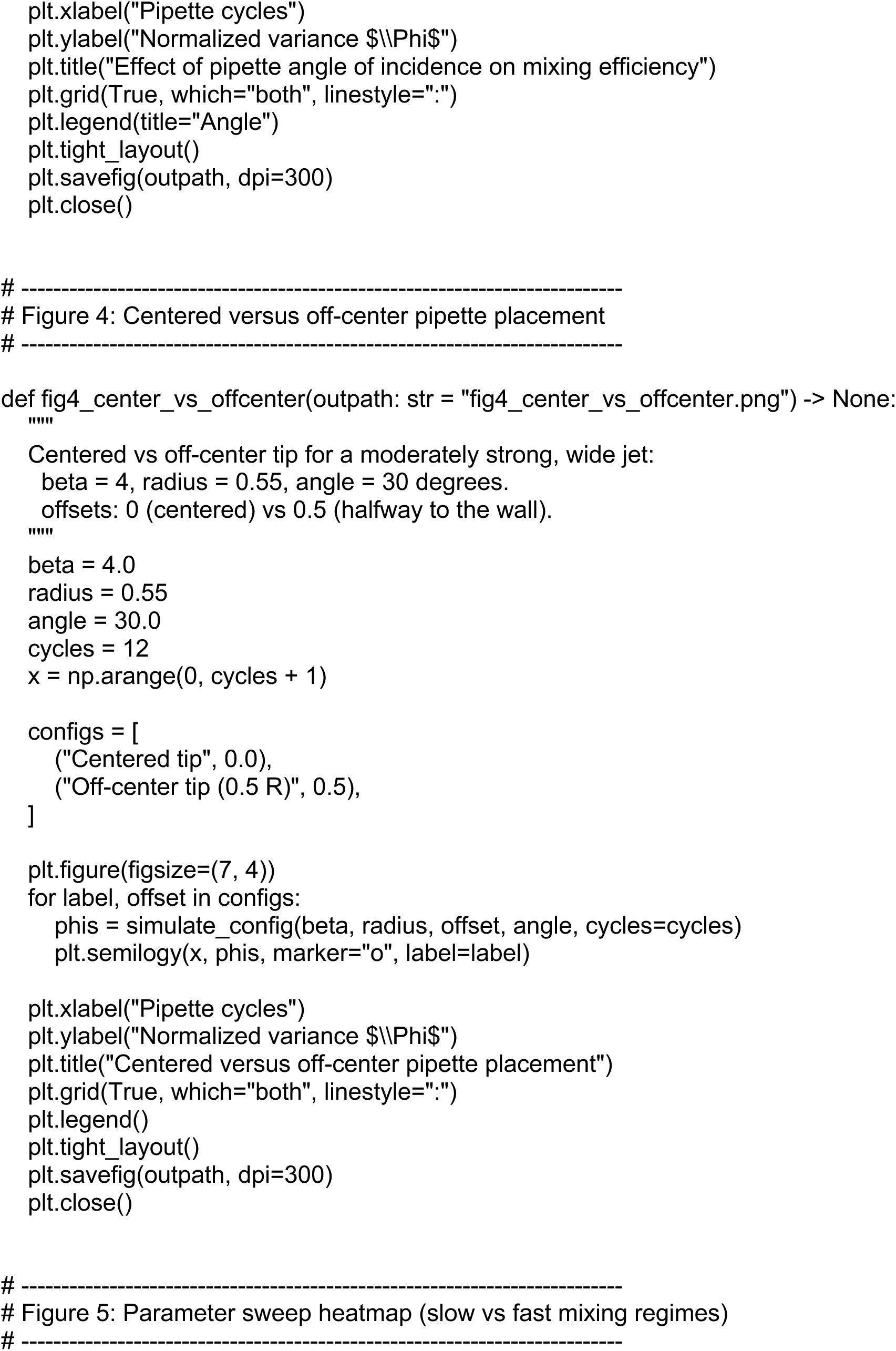
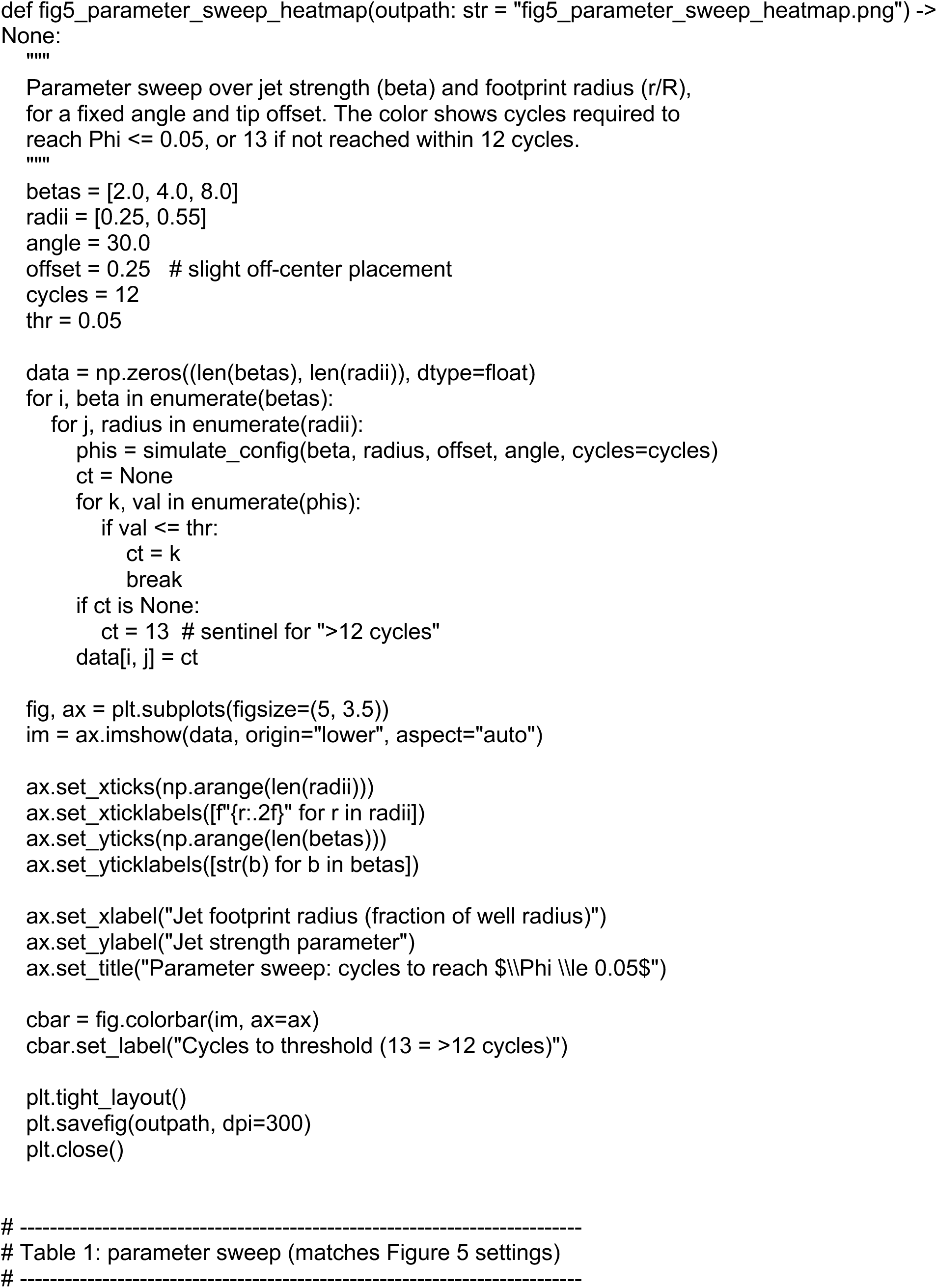
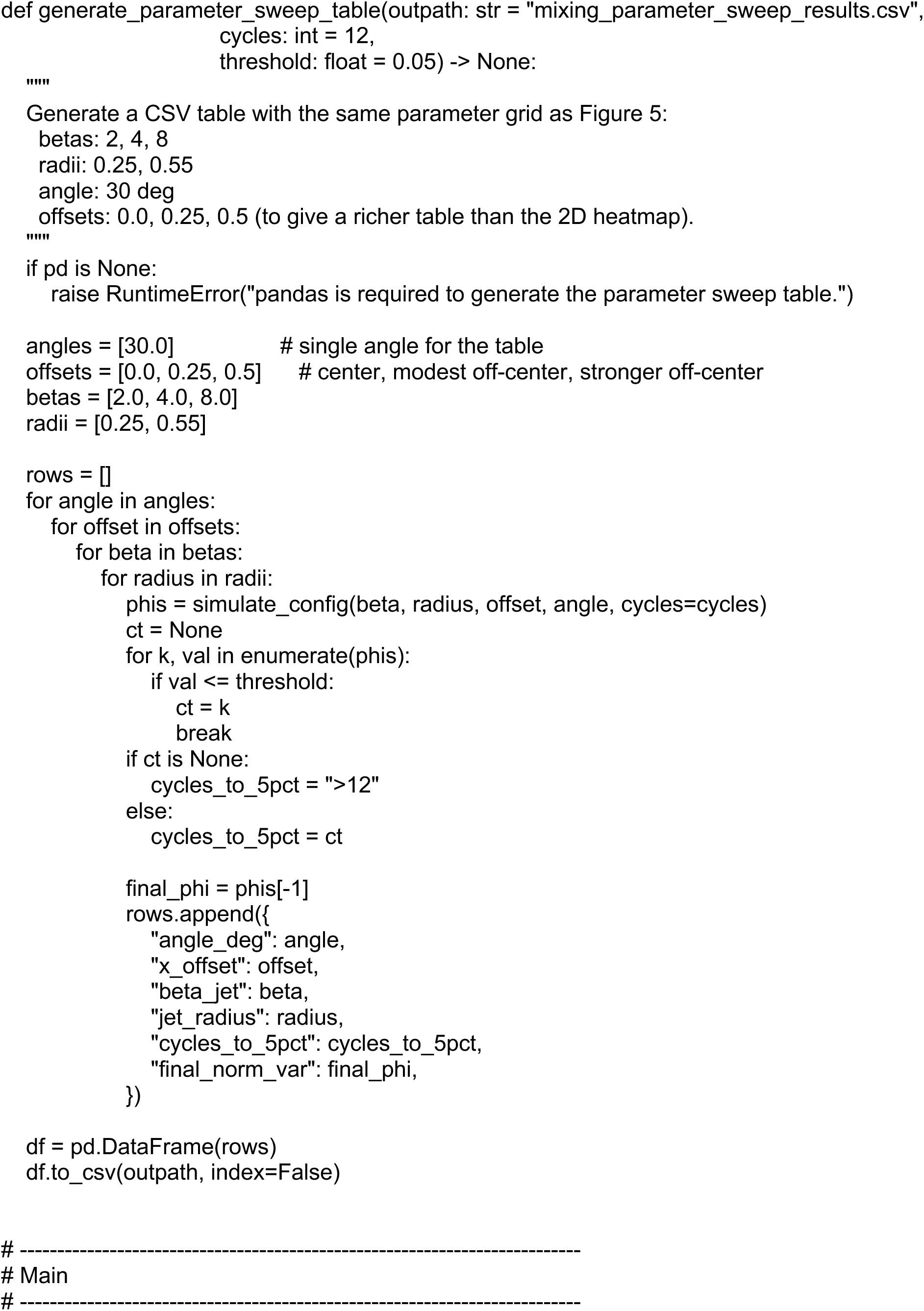
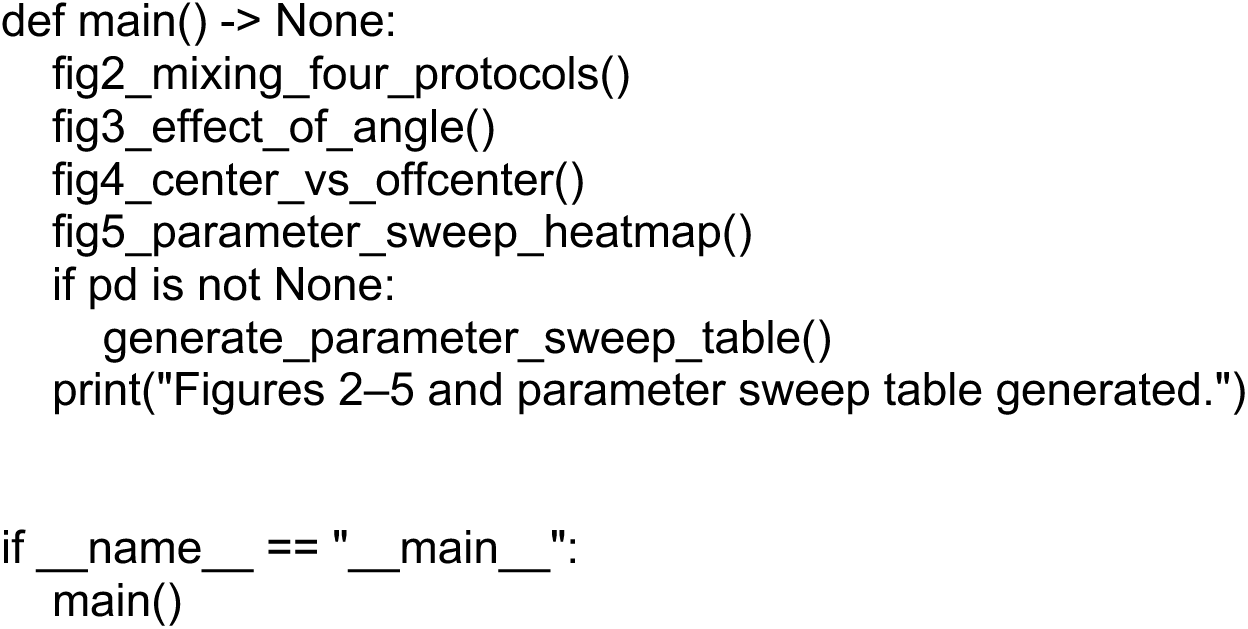

